# Mitotically Driven Cytoskeletal Reorganization Governs Zebrafish Left-Right Organizer Detachment from EVL and Lumen Morphogenesis

**DOI:** 10.64898/2026.03.18.712746

**Authors:** Yan Wu, Yiling Lan, Miriam Athena Allred, Carys Timpson, Heidi Hehnly

**Affiliations:** Syracuse University, Syracuse NY, 13210; Arizona State University, Tempe AZ, 85281; Cambridge University, Cambridge UK, CB2 1TN

## Abstract

Left–right asymmetry in vertebrate embryos is established by the left–right organizer (LRO), with the zebrafish Kupffer’s vesicle (KV) providing a tractable model for studying de novo epithelial morphogenesis. During KV formation, dorsal forerunner cells (DFCs) initially form polarized attachments to the enveloping layer (EVL) before reorganizing into multicellular rosettes that precede lumen formation. Here, we show that while DFC–EVL junctions form independently of mitosis, early cytokinetic events play an instructive role in remodeling these contacts. Live imaging and targeted laser ablation reveal that cytokinetic bridges and their associated microtubule bundles recruit actin, seed rosette centers, and promote the transition from external to internal epithelial organization. Disruption of early, but not later, DFC divisions impairs actin accumulation, rosette coalescence, KV detachment from the EVL, and lumenogenesis. These findings identify a temporally restricted role for cytokinesis in organizing cytoskeletal architecture and reveal how division history directs epithelial tissue assembly during LRO development.

## INTRODUCTION

During vertebrate development, establishment of the left–right axis depends on the formation and function of the left–right organizer (LRO) (Dasgupta and Amack, 2016; Grimes and Burdine, 2017; Lee and Anderson, 2008; Wells et al., 2022). In zebrafish the left-right organizer is known as the Kupffer’s vesicle (KV) (Essner et al., 2005), which provides a powerful in vivo system for dissecting the cellular and cytoskeletal mechanisms that underlie de novo organogenesis within the embryo. Although KV cilia-driven fluid flow has been extensively studied as a determinant of laterality (Dasgupta and Amack, 2016; Hirokawa et al., 2009; Kramer-Zucker et al., 2005; Okabe et al., 2008; Sampaio et al., 2014), far less is known about the cytoskeletal events that precede lumen formation and pattern the early architecture of the organ. How microtubule- and actin-based structures are coordinated to establish polarity, junctional organization, and tissue-level morphology during KV assembly remains incompletely understood. Notably, direct visualization of microtubule dynamics during early KV morphogenesis has only recently become feasible (Wu et al., 2025), enabling new insights into how microtubule organization contributes to the earliest stages of LRO assembly.

Prior studies of KV morphogenesis have largely emphasized actin-based mechanisms that shape the behavior of dorsal forerunner cells (DFCs), the progenitor population that gives rise to the cells that line the KV’s lumen. DFCs must undergo a series of coordinated morphogenic transitions: initially organizing into small, transient multicellular rosettes, subsequently coalescing into a larger rosette-like structure, and ultimately rearranging into a cyst of polarized epithelial-like cells surrounding a central lumen (Amack, 2021; Amack et al., 2007; Oteíza et al., 2008). Actomyosin contractility, regulated by RhoA signaling and associated guanine nucleotide exchange factors, has been shown to be essential for DFC cell shape changes, epithelialization, and rosette formation during these stages (Frisca et al., 2016; Wang et al., 2011; Wang et al., 2012; Zhang et al., 2016). Apical constriction and junctional remodeling have likewise been implicated in shaping the KV epithelium, and computational models have linked cell contractility, adhesion, and mechanical forces to rosette organization and lumen expansion (Dasgupta et al., 2018; Erdemci-Tandogan et al., 2018; Manna et al., 2025; Sanematsu et al., 2021). Together, these studies support a framework in which actin–myosin dynamics generate the forces required to drive DFC rearrangements and epithelial morphogenesis.

In contrast, relatively little is known about the specific contribution of cell–cell adhesion molecules, including E-cadherin, to KV morphogenesis. Classical genetic studies in zebrafish established that E-cadherin–mediated adhesion is essential for epithelial cohesion and coordinated morphogenetic movements during early embryogenesis. In a zebrafish epiboly mutant screen, the *half baked* locus was identified as encoding E-cadherin, and its loss disrupts blastoderm cell adhesion and collective cell movements during gastrulation and epiboly (Babb and Marrs, 2004; Kane et al., 1996; Shimizu et al., 2005). Despite this foundational role in early tissue organization, the function and regulation of E-cadherin during later morphogenetic transitions such as KV formation remain poorly defined. One study identified a genetic interaction between the polarity protein Lgl2 and the recycling endosome GTPase Rab11a in regulating E-cadherin localization at cell–cell contacts in a fully formed KV with lumen, while reporting comparatively modest effects on overall KV morphogenesis (Tay et al., 2013). However, how E-cadherin is organized and dynamically regulated during DFC rearrangements as they transition into a mature KV epithelium is unknown.

Building on this limited understanding of adherens junction dynamics, even less is known about the establishment and organization of tight junctions during KV morphogenesis. Tight junction components, including the scaffolding protein ZO-1, serve as key organizers of epithelial polarity and barrier formation in many developing tissues. As DFCs transition from loosely organized progenitors into a polarized epithelial cyst surrounding the nascent KV lumen, they need to establish apical junctional complexes that define tissue architecture and lumenal integrity. ZO-1 has been reported to localize to apical junctional regions of KV cells once a lumen is formed (Dasgupta et al., 2018; Lopes et al., 2010; Navis et al., 2013), suggesting that tight junction assembly accompanies epithelial maturation of the KV. However, when and how ZO-1–based junctions are first established, how they are coordinated with cytoskeletal remodeling, and whether their formation depends on earlier morphogenetic events such as cell division or rosette formation remain unresolved. These gaps highlight the need to define how tight junction scaffolds integrate with adhesion and cytoskeletal networks to support epithelial organization during KV development.

Insights from polarized epithelial systems indicate that cytokinesis can act as an instructive cue during epithelial morphogenesis. In cultured epithelial models, including MDCK cells, cytokinesis involves the coordinated formation of a new adhesive interface between daughter cells through cell rounding, membrane ingression, and de novo junction assembly. Seminal studies have shown that cytokinetic bridges and their associated midbodies function as spatial landmarks for junctional remodeling, actin polymerization, and apical membrane initiation, in part through the localized recruitment of trafficking machinery, cytoskeletal regulators, and polarity determinants (Addi et al., 2018; Dambournet et al., 2011; Dionne et al., 2015; Mangan et al., 2016; Schlüter et al., 2009; Sugioka, 2022; Wang et al., 2014). In vivo studies in Drosophila epithelia further demonstrate that cytokinetic actomyosin ring contraction is mechanically coupled to adherens junctions, triggering junctional remodeling in both dividing and neighboring cells through coordinated Rho GTPase signaling and myosin II dynamics (Daniel et al., 2018; di Pietro et al., 2023; Guillot and Lecuit, 2013; Herszterg et al., 2013; Mira-Osuna and Le Borgne, 2024; Pinheiro and Bellaïche, 2018; Pinheiro et al., 2017). Together, these studies establish cytokinesis as a potent organizer of epithelial architecture.

While these systems have provided critical insight into how cytokinesis interfaces with epithelial architecture, they do not directly address how similar mechanisms operate during developmental contexts in which epithelial polarity emerges progressively rather than being pre-established. Evidence from developmental systems supports this possibility: in the early *C. elegans* embryo, the LKB1 ortholog PAR-4 coordinates actomyosin contractility during both polarity establishment and cytokinesis, with defects in cytokinetic dynamics leading to mispositioned polarity determinants (Chartier et al., 2011). Consistent with this idea, our previous work in zebrafish showed that cytokinetic bridges marked by MKLP1 are highly ordered during KV formation, orient toward rosette centers, and require timely abscission for lumen opening (Rathbun et al., 2020). These observations suggest that cytokinesis-associated structures can act as transient but instructive organizers of cytoskeletal and junctional remodeling during de novo epithelial organogenesis.

KV development presents a powerful in vivo context in which to examine how cytokinesis interfaces with epithelial organization as polarity emerges. DFCs initially maintain polarized attachments to the enveloping layer (EVL) (Hirokawa et al., 2006; Hirokawa et al., 2009; Oteíza et al., 2008). These attachments serve as organizational “seeds” for KV formation but must be extensively remodeled as DFCs proliferate, reorganize into multicellular rosettes, and ultimately assemble a lumenized epithelium. Notably, DFCs undergo multiple rounds of cell division during this transition, and our recent work demonstrated that the earliest mitotic events are essential for timely KV lumen formation, whereas later divisions are largely dispensable (Wu et al., 2025). The cellular mechanisms underlying this stage-specific requirement, however, have remained unresolved.

Here, we sought to define how E-cadherin–mediated adhesion, tight junction scaffolding, and cytoskeletal remodeling coordinate DFC morphogenesis into a functional KV. Using endogenously tagged E-cadherin and ZO-1 zebrafish lines, we performed high-resolution live imaging from shield stage through the 3-somite stage, capturing the transition from a loosely associated DFC cluster to a lumenized epithelial cyst. We find that ZO-1 dynamically relocalizes in concert with actin remodeling as DFCs detach from the EVL and reorganize into rosette structures, whereas E-cadherin remains stably maintained at lateral cell–cell interfaces and does not enrich at actin-dense rosette centers. Actin redistribution from DFC–EVL attachments to rosette foci is cell division–dependent, with cytokinetic microtubule structures promoting localized actin recruitment and rosette assembly. Collectively, these findings support a model in which stable cadherin-based adhesion provides structural continuity during DFC rearrangement, while tight junction scaffolding and cytokinesis-driven cytoskeletal remodeling actively drive epithelialization and KV lumen formation.

## RESULTS

### E-cadherin remains localized at DFC cell–cell Interfaces during actin remodeling in KV morphogenesis

To determine how cadherin-based adhesion behaves during the dynamic cellular rearrangements that accompany KV morphogenesis, we sought to establish the three-dimensional organization of DFC populations at defined developmental transitions. DFCs undergo detachment from the EVL and compaction into a rosette-like intermediate prior to lumen formation (Figure 1A). These events are accompanied by pronounced actin remodeling (Amack et al., 2007), raising the question of whether E-cadherin–mediated cell–cell adhesion is similarly redistributed or instead remains stably maintained as tissue architecture is reconfigured. To directly assess the spatial relationship between E-cadherin and actin during KV morphogenesis, we performed high-resolution 3D confocal imaging of embryos carrying an endogenously tagged *cdh1*-mlanYFP (E-cadherin) allele (Cronan and Tobin, 2019), fixed and stained for F-actin (Figure 1B and 1C). Embryos were collected at 8 (Figure 1B) and 12 hpf (Figure 1C)—stages corresponding to early DFC clustering/rosette formation and established lumen formation, respectively—and the KV volume was reconstructed in three dimensions relative to the overlying EVL (Figure 1B and 1C). These reconstructions enabled quantitative visualization of cell–cell interfaces within both the surface epithelium and the underlying DFC/KV population in intact embryos. At the level of the EVL, E-cadherin localization closely paralleled cortical actin organization, consistent with canonical adherens junction architecture. In contrast, within the DFC/KV compartment, E-cadherin distribution did not mirror actin reorganization. Optical sections through KV cells at 8 hpf revealed emerging rosette structures characterized by focal actin enrichment at their centers (Figure 1D). While E-cadherin was robustly maintained along lateral cell–cell interfaces, it was not enriched at actin-dense rosette foci (Figure 1D, arrow). Line scan analysis across these interfaces confirmed this observation: prominent peaks of actin intensity at rosette centers lacked a corresponding increase in E-cadherin signal (Figure 1F).

**Figure 1.**
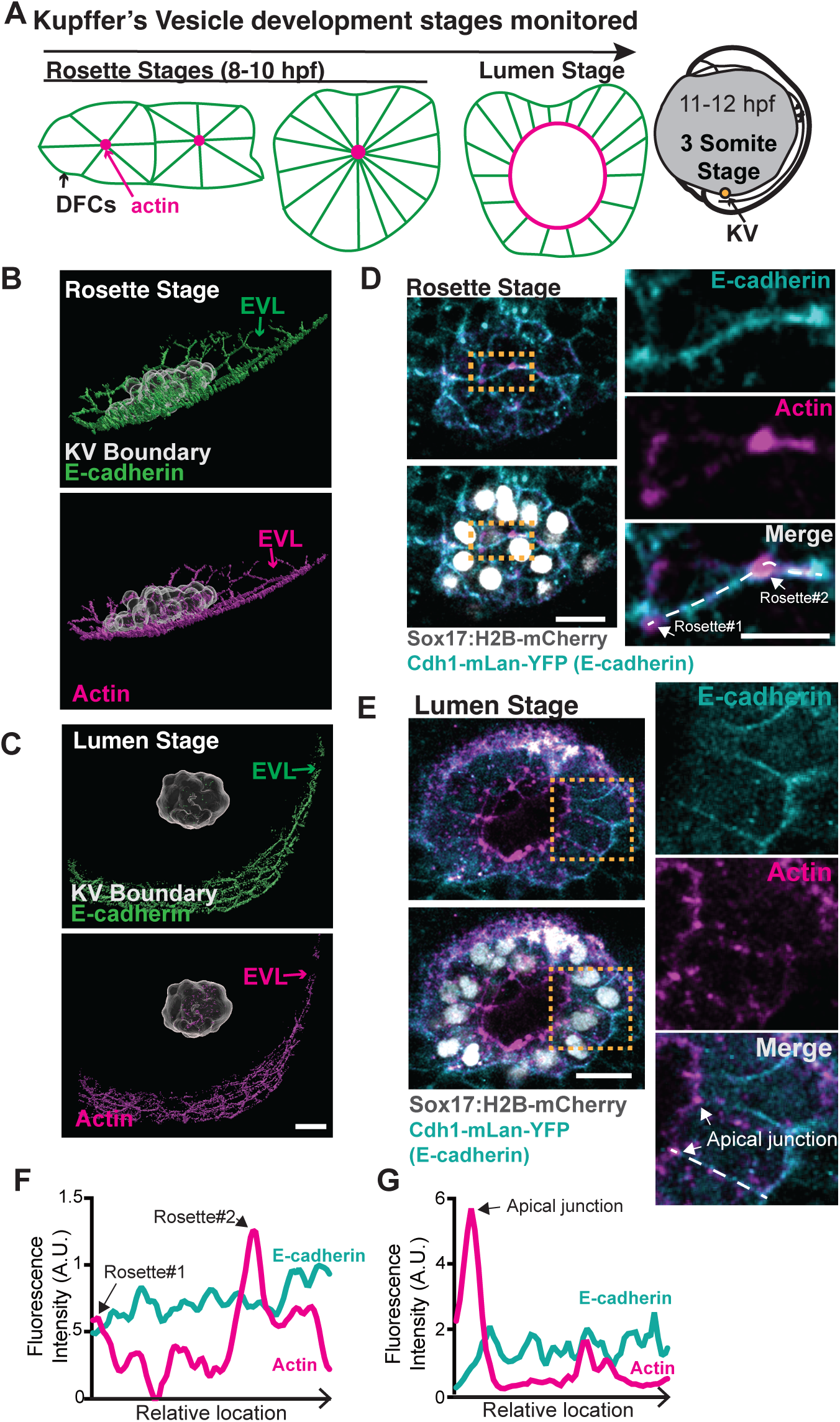
E-cadherin remains localized at DFC cell–cell Interfaces during actin remodeling in KV morphogenesis. **A)** Model depicting the represented developmental stages of KV in zebrafish embryos from 8 to 12 hours post-fertilization (hpf), including rosette and lumen stages. **(B-C)** 3D surface renderings generated from confocal z-stacks of representative embryos at (8 hpf, Rosette stage, **B**; and 12 hpf, Lumen stage, **C**), showing E-cadherin (green) and F-actin (phalloidin staining, magenta) puncta in dorsal forerunner cells (DFCs, gray) of Sox17:H2B-mCherry; Cdh1-mLan-YFP embryos. The enveloping layer (EVL) is marked. KV cells are marked by Sox17:H2B-mCherry. Scale bar, 30 μm. **(D–E)** Confocal maximum intensity projections of E-cadherin and F-actin puncta in the KV at the rosette stage (**D**) and Lumen stage (**E**) in Sox17:H2B-mCherry; Cdh1-mLan-YFP embryos. E-cadherin is shown in cyan (Cdh1-mLan-YFP), nuclei in white (Sox17:H2B-mCherry), and F-actin in magenta (phalloidin staining). Scale bars, 20 μm; Magnified image, 10 μm. **(F-G)** Fluorescence intensity profiles of E-cadherin and actin along the white dotted lines indicated in panels **D** and **E**.

By 12 hpf, when KV cells had organized into a cyst surrounding a centralized lumen, actin became enriched at the apical surface adjacent to the nascent lumen, coincident with the region where tight junction markers such as ZO-1 are localized (Figure 1E, arrow; (Dasgupta et al., 2018; Lopes et al., 2010; Navis et al., 2013)). Despite this apical actin accumulation, E-cadherin remained uniformly distributed along lateral cell–cell interfaces and did not concentrate at the apical domain (Figure 1E). Line scan quantification demonstrated a clear spatial separation between apical actin enrichment and E-cadherin localization (Figure 1G). This pattern is consistent with prior immunofluorescence analyses (Tay et al., 2013) and mirrors the distribution observed during the earlier rosette stage.

These findings suggest that although E-cadherin remains stably maintained at DFC/KV cell–cell contacts throughout morphogenesis, its distribution does not track with actin reorganization during rosette assembly or lumen formation. In contrast to the EVL, where E-cadherin and actin are spatially coordinated, actin reorganization within the DFC/KV population occurs without corresponding redistribution of E-cadherin. These observations indicate that dynamic actin enrichment at rosette centers and apical domains is not accompanied by local accumulation of E-cadherin.

### DFCs first attach to the EVL through ZO-1 and actin, then detach during rosette formation, KV rounding, and lumenogenesis

Our initial analyses revealed that, unlike in the EVL, E-cadherin does not distribute with actin during rosette assembly or apical remodeling within the forming KV. This raised the question of which junctional architecture is coordinated with the pronounced actin reorganization that accompanies DFC detachment and epithelialization. Prior studies established that DFCs arise through Nodal/TGFβ-dependent ingression from the dorsal surface epithelium and that a subset of these cells retains apical attachments to the overlying EVL (Oteíza et al., 2008). These persistent contacts were proposed to act as polarity “seeds,” transmitting epithelial organization from the EVL to the internal KV epithelium. However, the molecular composition of these interfaces and how they are dynamically remodeled during rosette formation and lumenogenesis remain incompletely defined.

To define how tight junction scaffolds and associated apical actin networks are organized during DFC cellular rearrangements underlying KV morphogenesis, we performed high-resolution 3D confocal imaging and volumetric reconstruction of intact embryos fixed at 6, 9, 10, and 12 hpf, spanning stages from early attachment through lumen establishment (Figure 2A). DFCs were selectively labeled using the transgenic line *Sox17:EMTB-3xGFP*, enabling identification and three-dimensional segmentation of KV-fated cells within confocal datasets (Figure 2B, 2C, Video S1). EVL cells were independently identified by cortical actin staining, and embryos were immunostained for the tight junction marker ZO-1 to resolve junctional assemblies.

**Figure 2.**
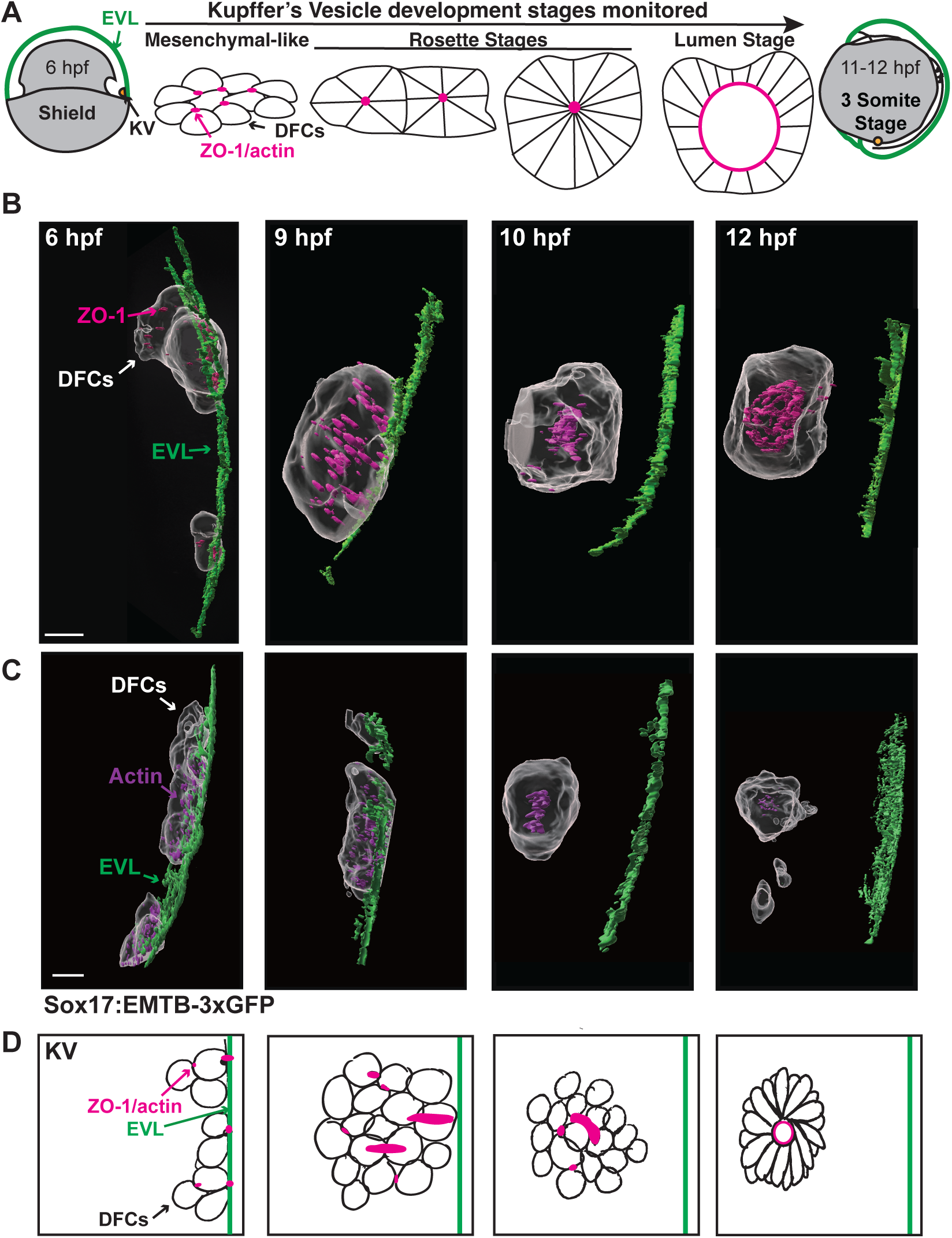
DFCs first attach to the EVL through ZO-1 and actin, then detach during rosette formation, KV rounding, and lumenogenesis. **(A)** Top-down model depicting the developmental stages of Kupffer’s Vesicle (KV) in zebrafish embryos from 6 to 12 hpf, including mesenchymal-like, rosette, and lumen stages. **(B)** 3D surface renderings of representative embryos showing dorsal forerunner cells (DFCs, gray) detaching from the enveloping layer (EVL, green), with ZO-1 visualized in magenta. The KV boundary is marked using Sox17:EMTB-3xGFP. Scale bar: 20 µm. Refer to Supplemental Video 1. **(C)** 3D renderings highlighting actin (purple). Scale bar: 20 µm. **(D)** A side-view models depicting how DFCs initially attach to the EVL through ZO-1 and actin, then detach during rosette formation, KV rounding, and lumenogenesis.

At early stages, side-view reorientation of reconstructed 3D volumes revealed a similar distribution pattern of ZO-1 (Figure 2B) and actin (Figure 2C) at DFC–EVL interfaces, delineating discrete attachment zones along the DFC border consistent with persistent epithelial contacts. By 9 hpf, as the KV began to round, this organization shifted: ZO-1– and actin-positive domains became progressively enriched at DFC–DFC interfaces oriented toward the interior of the structure. Although residual DFC–EVL attachments remained detectable, the overall architecture indicated a transition from external anchoring to internally focused junctional assemblies, consistent with the emergence of overlapping rosette-like structures within the DFC cluster.

By 10 hpf, the KV had fully detached from the EVL, coincident with consolidation of a centralized rosette architecture composed exclusively of DFC–DFC interactions. At this stage, ZO-1 and actin were no longer detectable at DFC–EVL interfaces and were instead confined to junctional domains within the DFC population. By 12 hpf, as the KV internalized further and initiated lumen formation, ZO-1 and actin became highly organized at the apical surfaces of DFCs lining the nascent lumen, marking the establishment of polarized epithelial architecture. Together, these observations define a stepwise redistribution of actin- and ZO-1–positive junctions: from EVL-anchored attachment sites to internally oriented rosette interfaces, and finally to apical domains surrounding the forming lumen (side view oriented model, Figure 2D).

### ZO-1 site assembly precedes actin; ZO-1 and actin then accumulate together at DFC-EVL interfaces

Our analysis in Figure 2 revealed that sites of contact between DFCs and the EVL are characterized by the co-enrichment of the tight junction protein ZO-1 and filamentous actin. These observations indicate that DFC–EVL interfaces adopt molecular features of epithelial junctions early in KV morphogenesis, even when DFCs themselves exhibit “mesenchymal-like” organization (Amack, 2021; Amack et al., 2007). However, the close spatial coupling of ZO-1 and actin at these sites does not resolve the order of assembly of junctional and cytoskeletal components. Determining whether junctional scaffolds are established prior to actin recruitment, or conversely whether actin accumulation precedes and stabilizes ZO-1–based contacts, is essential for understanding how transient DFC–EVL attachments are initiated.

To address this question, we examined the earliest stages of DFC–EVL contact formation, focusing on 6-7 hpf embryos when DFCs are dispersed and have not yet organized into multicellular rosettes. DFCs were unambiguously distinguished from EVL cells using Sox17-driven reporters, either Sox17:EMTB-3xGFP (Figure 3A–D; EMTB signal not shown for clarity) or Sox17:H2B-mCherry (Figure 3E–F; visible in Video S2), enabling cell-type–specific identification throughout the analysis. In fixed embryos (Figure 3A–D), we simultaneously immunostained for ZO-1 and labeled filamentous actin using phalloidin to assess their relative spatial organization at nascent contact sites. In parallel, we employed a CRISPR-engineered zebrafish line in which endogenous ZO-1 (tjp1) is fluorescently tagged with tdTomato (Levic et al., 2021), enabling live visualization of ZO-1 dynamics under physiological expression levels (Figure 3E-F, Video S2). In these embryos, actin organization was monitored using Lifeact-mEmerald, allowing direct, time-resolved comparison of ZO-1 and actin recruitment at DFC–EVL interfaces. Together, these complementary fixed and live imaging approaches were designed to define the temporal hierarchy of junctional and cytoskeletal assembly underlying DFC–EVL adhesion.

**Figure 3.**
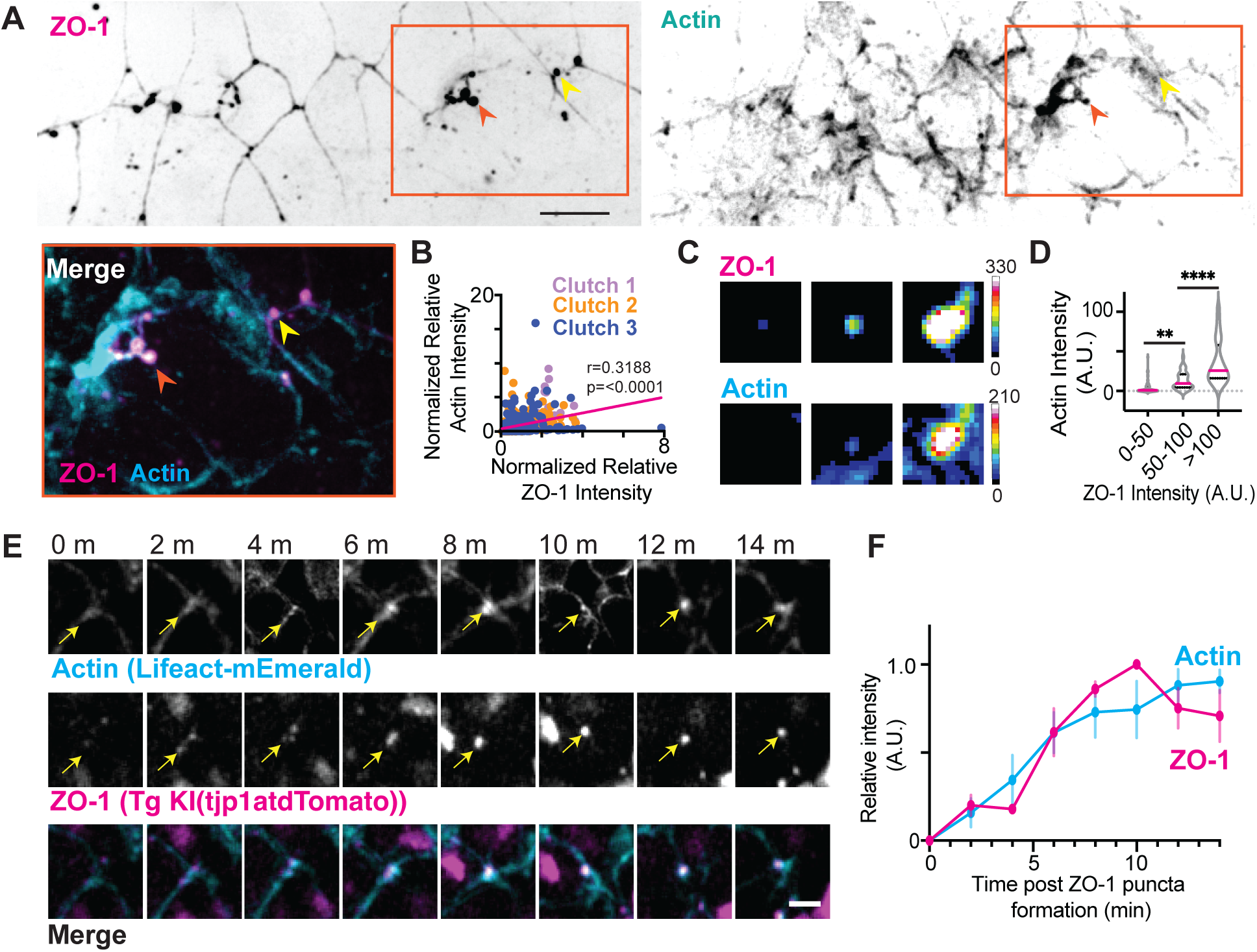
ZO-1 site assembly precedes actin; ZO-1 and actin then accumulate together at DFC-EVL interfaces. **(A)** Representative images of ZO-1 (left; magenta in merge) and actin (right; cyan in merge) at DFC-EVL interface at 7 hpf. A large ZO-1 punctum with actin recruitment is indicated by an orange arrow, and a small ZO-1 punctum lacking actin recruitment is indicated by a yellow arrow. The merged region outlined by the orange box is shown at higher magnification below. Scale bar: 20 μm. **(B)** Scatter plot showing a positive correlation between normalized actin intensity and normalized ZO-1 intensity at ZO-1 puncta across three clutches (light purple, orange, blue). The magenta line indicates the linear trend where slope=r. Pearson’s r = 0.3188, P < 0.0001. (3 clutches; 118 embryos; 494 puncta)**. (C)** Representative images of examples of puncta with ZO-1 intensities between 0-50 A.U. (left), 50-100 A.U. (middle), and >100 A.U (right). ZO-1 and actin intensities are displayed above their respective images. Calibration bars showing the range of intensities for both ZO-1 and Actin are shown to the right of the images. **(D)** A representative Violin plot from 1 clutch of actin intensity at ZO-1 puncta binned by ZO-1 intensity ranges (0–50, 50–100, >100 A.U.). Mean values are indicated by magenta lines and quartiles by black lines. **P < 0.01, ****P < 0.0001 (one-way ANOVA with Dunnett’s multiple comparisons; n = 28 embryos, 131 puncta). **(E)** Time-lapse images showing actin (Lifeact-mEmerald, top; cyan in merge) and ZO-1 (middle; magenta in merge) recruitment at KV puncta. Scale bar: 10 μm. **(F)** Relative intensity of actin and ZO-1 at puncta in KV over time across n=3 puncta followed from 1 live embryo. Error bars represent s.e.m. Refer to Supplemental Video 2 and Supplemental Table S1.

In early embryos containing fewer than 20 KV cells, ZO-1 was detected as small, discrete puncta at DFC–EVL interfaces, typically on the order of ∼1 μm² in area. At this stage, many ZO-1 puncta exhibited little to no associated actin enrichment (Figure 3A, yellow arrow; Figure 3C). A subset of puncta, however, was larger and displayed higher ZO-1 fluorescence intensity (Figure 3A, orange arrow; Figure 3C), suggestive of more mature junctional assemblies. To assess the relationship between ZO-1 accumulation and actin recruitment, we quantified the integrated fluorescence intensity of individual ZO-1 puncta and extracted the corresponding phalloidin signal from the same segmented regions; values were normalized to the mean puncta intensity within each embryo (Figure 3B). Normalized actin intensity exhibited a modest but significant positive correlation with ZO-1 intensity across puncta (r = 0.3188, p < 0.0001). This trend was consistent across clutches, indicating that increased ZO-1 accumulation is associated with enhanced local actin enrichment. However, the relatively low correlation coefficient suggests substantial variability at the level of individual puncta, which may reflect biological or clutch-to-clutch variation. To further examine this relationship without normalization to the clutches mean intensity, we analyzed a representative clutch and found that actin intensity remained low at ZO-1 puncta with weak fluorescence (0–50 A.U.), but increased once ZO-1 intensity exceeded a threshold of ∼50 A.U. (Figure 3C–D). Together, these fixed-embryo analyses support a model in which ZO-1 site assembly and maturation precede robust actin recruitment at DFC–EVL contacts.

To directly examine the dynamics underlying this transition, we next performed live imaging of ZO-1 and actin during early KV development using embryos expressing endogenously tagged ZO-1 (Tjp1–tdTomato) together with Lifeact-mEmerald. In contrast to fixed samples, we rarely observed ZO-1 puncta entirely devoid of actin in live embryos (Figure 3A, 3C, compared to 3E). This likely reflects differences in detection sensitivity, as endogenous ZO-1–tdTomato signal was comparatively dim relative to the high expression levels of Lifeact driven by injected mRNA. Despite this limitation, time-resolved imaging revealed that nascent ZO-1 puncta initially appeared as small, low-intensity assemblies that progressively increased in size and fluorescence intensity over 15 minutes (Figure 3E-F). Importantly, actin intensity at these sites increased in parallel with ZO-1 growth, exhibiting coordinated accumulation kinetics rather than delayed recruitment.

Together, the fixed and live imaging data support a model in which ZO-1–based junctional sites are initiated first at DFC–EVL interfaces, reaching a critical size or molecular density that stabilizes subsequent actin enrichment. Live imaging further suggests that, once initiated, ZO-1 and actin co-accumulate in a tightly coupled manner as junctions mature. This stepwise assembly provides a mechanistic framework for how transient epithelial attachments between DFCs and the EVL are established.

### Following DFC-EVL attachments, DFC cytokinetic events converge and precede rosette formation

Our findings thus far suggest that DFC–EVL attachments are initiated through ZO-1–based junctional assemblies that recruit actin, generating polarized, epithelial-like contact sites. As KV development proceeds, DFCs undergo an expansion in cell number (Rathbun et al., 2020; Wu et al., 2025) accompanied by a reorganization of cell–cell interactions, transitioning from an EVL-attached population into multicellular rosette structures (Figure 2A). In the context of DFCs, a rosette is defined as a radially organized assembly in which multiple cells converge around a shared focal point enriched in apical junctional components (e.g., ZO-1; Figure 2A) and cytoskeletal elements (e.g., actin; Figure 2A). This configuration represents an intermediate architectural state that precedes formation of a single, centralized epithelial lumen (Gredler and Zallen, 2023; Harding et al., 2014). In the case of KV development, it is unclear how the actin-enriched DFC–EVL attachments characterized in Figures 2 and 3 are remodeled as DFCs begin to proliferate at ∼6-8 hpf, and whether cell division plays an instructive role in organizing rosette architecture. DFC proliferation coincides temporally with the earliest appearance of rosette-like assemblies (Krishnan et al., 2022; Rathbun et al., 2020; Wu et al., 2025), raising the possibility that cytokinetic events may actively contribute to rosette initiation rather than occurring as a downstream consequence of epithelialization. Cytokinesis is known to locally reorganize actin and microtubules and to generate transient intercellular connections, features that could provide spatial cues for the convergence of neighboring cells (Mira-Osuna and Le Borgne, 2024). Thus, determining whether DFC divisions precede, coincide with, or follow rosette formation is critical for understanding how early junctional contacts are repurposed to drive higher-order tissue organization.

We therefore asked whether the timing and positioning of DFC divisions are linked to actin-rich contact sites and the emergence of rosette centers, thereby coupling proliferative dynamics to cytoskeletal remodeling during early KV morphogenesis. To address this, we performed live imaging in embryos expressing Sox17:EMTB-3xGFP (used in (Wu et al., 2025)) to visualize DFC microtubules and cytokinetic bridges, together with Lifeact-mRuby to monitor actin organization. Imaging was initiated at 6 hpf, prior to overt rosette formation, enabling individual cytokinetic events to be tracked with high temporal resolution.

In Figure 4A, we tracked a single DFC division over a 42-minute interval. At the onset of imaging (0 min), the dividing cell was in metaphase, with a clearly defined mitotic spindle oriented along the longest axis of the KV. Notably, this DFC maintained two discrete junctional contact sites with an overlying EVL cell. As division progressed, these DFC–EVL contact sites did not dissipate but instead moved toward one another, converging toward the site of the forming cytokinetic bridge. By 16 minutes, the former attachment sites were spatially reorganized around the midbody region, and by 42 minutes, this convergence coincided with the emergence of a nascent rosette center. Importantly, no pre-existing rosette was present prior to this division event, indicating that cytokinesis precedes and potentially predicts the site of rosette initiation.

**Figure 4.**
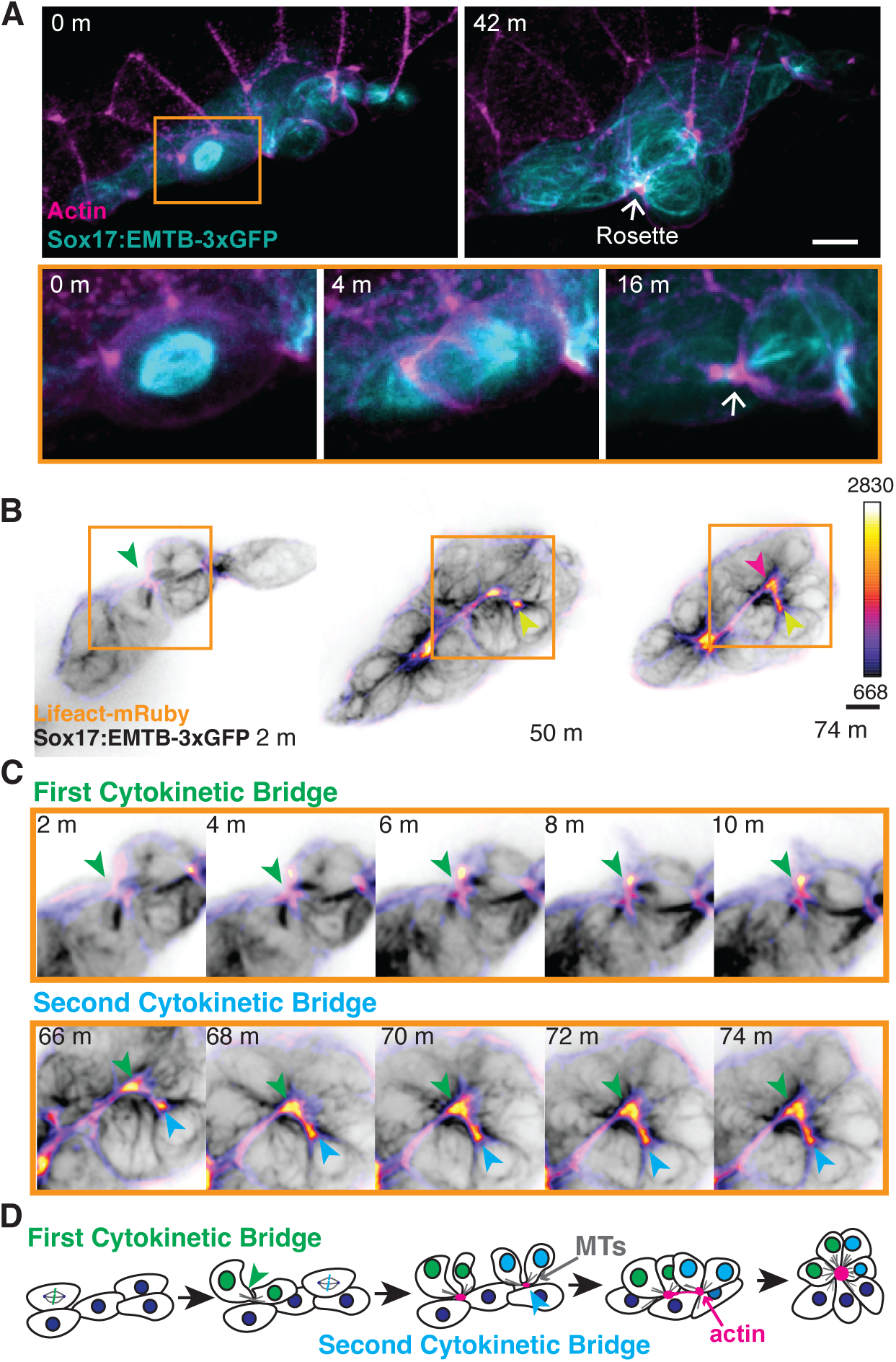
Following DFC-EVL attachments, DFC cytokinetic events converge and precede rosette formation. **(A)** Representative images from timeseries showing actin (Lifeact-mRuby, magenta) organization with a dividing cell and ultimately a cytokinetic bridge (microtubules, cyan) assembling at a forming rosette in the KV. 2.5x magnification shown for mitotic cell at 0 minute (m), 4 m, and 16 m. Scale bar: 10 μm. Microtubules labeled in KV with Sox17:EMTB-3xGFP embryos. **(B-C)** Time-lapse imaging of actin (Lifeact-mRuby, FireLUT) and microtubules (Sox17:EMTB-3xGFP, inverted gray). The region outlined by the orange box in (**B**) is shown at 1.5X magnification in (**C**). The green arrow highlights the first cytokinetic bridge relative to the second bridge (blue arrow), converging at the rosette. Scale bar: 10 μm. Calibration bar indicates Lifeact-mRuby fluorescence intensity. Refer to Supplemental Video 3. **(D)** Proposed temporal and spatial model illustrating how cytokinetic bridge–derived microtubule (MT) bundles recruit actin to seed initial rosette formation within KV.

We next following multiple division events over longer time scales (Figure 4B–C). In these recordings, the first cytokinetic event generated a stable cytokinetic bridge that marked the site of a forming rosette, accompanied by progressive actin accumulation at this focal point (green arrow, 2–10 min, Figure 4C). A second DFC division subsequently occurred within the vicinity of this established rosette center (blue arrow, inset, 66–74 min, Figure 4C, Video S3), with its cytokinetic bridge converging toward the same actin-rich site. Actin signal at the rosette center continued to intensify following the second division, suggesting that successive cytokinetic events reinforce and stabilize the emerging rosette architecture rather than disrupting it. The model in Figure 4D summarizes this process, illustrating how sequential DFC cytokinetic events converge on a shared focal point to nucleate multicellular rosette structures. These findings support a framework in which proliferative dynamics play an instructive role in early KV morphogenesis.

### Cytokinetic bridge derived microtubule bundles facilitate actin organization

The live-imaging analyses in Figure 4 revealed a strong spatial and temporal correlation between DFC cytokinetic bridges and the emergence of actin-enriched rosette centers. However, whether cytokinetic bridges play an instructive role in recruiting actin, or instead merely coincide with sites of actin accumulation, remain unresolved. We therefore examined whether intact cytokinetic bridges are required for actin recruitment during the earliest stages of rosette formation. To test this we performed targeted laser ablation experiments in Sox17:EMTB-3xGFP embryos injected with Lifeact-mRuby, allowing simultaneous visualization of cytokinetic microtubule bundles and actin dynamics. Experiments were conducted at 6 hpf, prior to the appearance of multicellular rosettes, when cytokinetic bridges can be unambiguously identified. Using a 355 nm laser, short line ablations (∼2 μm) were applied to cytokinetic microtubule bundles before robust actin accumulation at the bridge. This approach enabled selective disruption of cytokinetic architecture while minimizing collateral damage to surrounding cells (Figure 5A, B).

**Figure 5:**
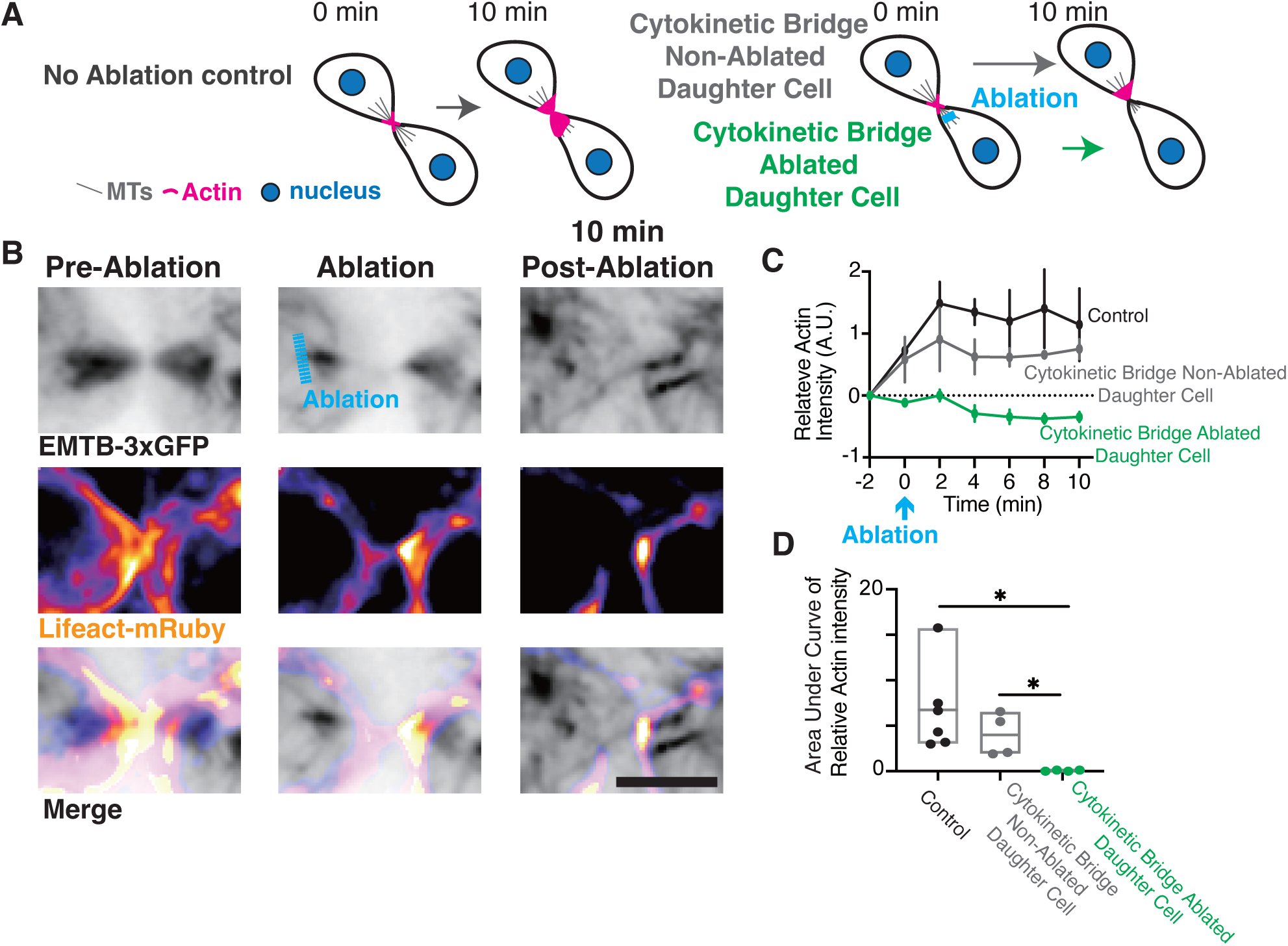
Cytokinetic bridge derived microtubule bundles facilitate actin organization. **(A)** Model illustrating actin dynamics of the daughter cells still connected by a cytokinetic bridge either without laser ablation (left) or with targeted ablation (right). In the ablation condition, one daughter cell is exposed to laser ablation at the bridge (blue), distinguishing between the non-ablated and ablated daughter cells. **(B)** Time-lapse images of microtubules (MTs) (Sox17:EMTB-3xGFP, Grey) and actin (Lifeact-mRuby, FireLUT) organization during pre-ablation, ablation and 10 minutes post-ablation. Microtubule and actin dynamics are disrupted following ablation. Scale bar: 10 µm. **(C)** Quantification of relative actin intensity over time for control (black), non-ablated daughter cells (gray), and ablated daughter cells (green), showing a significant reduction in actin accumulation after ablation. *n*≥4 embryos for each group. **(D)** Area under the curve (AUC) analysis of actin intensity from panel **(C)**, confirming significantly reduced actin accumulation in ablated daughter cells. Data are represented as mean ± SD. *p < 0.05 (unpaired t-test). Refer to Supplemental Table S1.

To isolate the contribution of the cytokinetic bridge to actin recruitment, we compared three conditions (Figure 5A): (i) non-ablated controls, (ii) partial bridge ablation in which microtubule bundles associated with one daughter cell were severed while the opposing daughter cell remained intact, and (iii) the unablated sister daughter cell within the same division, serving as an internal control. Actin recruitment was quantified by measuring Lifeact-mRuby intensity at the cytokinetic bridge region over a 10-minute time window following ablation (Figure 5C). Daughter cells whose cytokinetic microtubule bundles were ablated exhibited a pronounced defect in actin accumulation at the bridge compared to both non-ablated controls and their intact sister cells (Figure 5C–D). Quantitative comparison using area-under-the-curve analysis of relative actin intensity time courses revealed a significant reduction in actin recruitment specifically in the ablated condition. These results suggest that cytokinetic bridges are not merely correlated with actin enrichment but potentially are required for efficient actin recruitment at nascent rosette sites.

### Early mitotic events drive DFC rosette transition and detachment from EVL

Our previous work demonstrated that the earliest DFC divisions play a disproportionate role in KV morphogenesis, as selective ablation of the first several mitotic events, when fewer than 20 DFCs are present, resulted in severe delays and defects in KV lumen formation, whereas ablation of later divisions had minimal impact (Wu et al., 2025). Although these findings established a temporal requirement for early mitotic events, the cellular mechanisms underlying this requirement remained unclear. Considering our current data showing that cytokinetic bridges may recruit actin and potentially seed rosette formation (Figure 4–5), we hypothesized that early DFC divisions drive a critical cytoskeletal reorganization that promotes rosette assembly and facilitates the transition from DFC–EVL to DFC–DFC attachments. In this model, mitotic events are not required to initiate DFC–EVL contacts per se, but are essential for reorganizing these attachments into internal, actin-rich rosettes that precede KV detachment from the EVL and subsequent lumenogenesis.

To test this model, we performed targeted ablation of early versus late DFC mitotic events in Sox17:EMTB-3xGFP embryos, following an experimental paradigm similar to that used in our prior study ((Wu et al., 2025), Figure 6A). Early mitotic ablations were performed at ∼6 hpf, when fewer than 20 DFCs are present, whereas late mitotic ablations were carried out after DFC number expanded beyond 20, a time point when we start to note small rosettes forming. Embryos were fixed at this point (∼7.5-8 hpf) and immunostained for ZO-1 to label junctional complexes, while actin staining delineated the EVL. We assessed whether mitotic ablation perturbed DFC–EVL cell–cell contacts. Using volumetric segmentation approaches analogous to those employed in Figure 1 and 2, DFCs were reconstructed relative to the EVL, and ZO-1 puncta were classified as residing at DFC–EVL interfaces or at DFC–DFC interfaces. Quantitative analysis revealed no significant differences in either the number or area of ZO-1 puncta under early or late mitotic ablation conditions compared to non-ablated controls (Figure 6B–D). These results indicate that mitotic ablation does not disrupt the formation or maintenance of DFC–EVL attachments, consistent with our proposed model.

**Figure 6:**
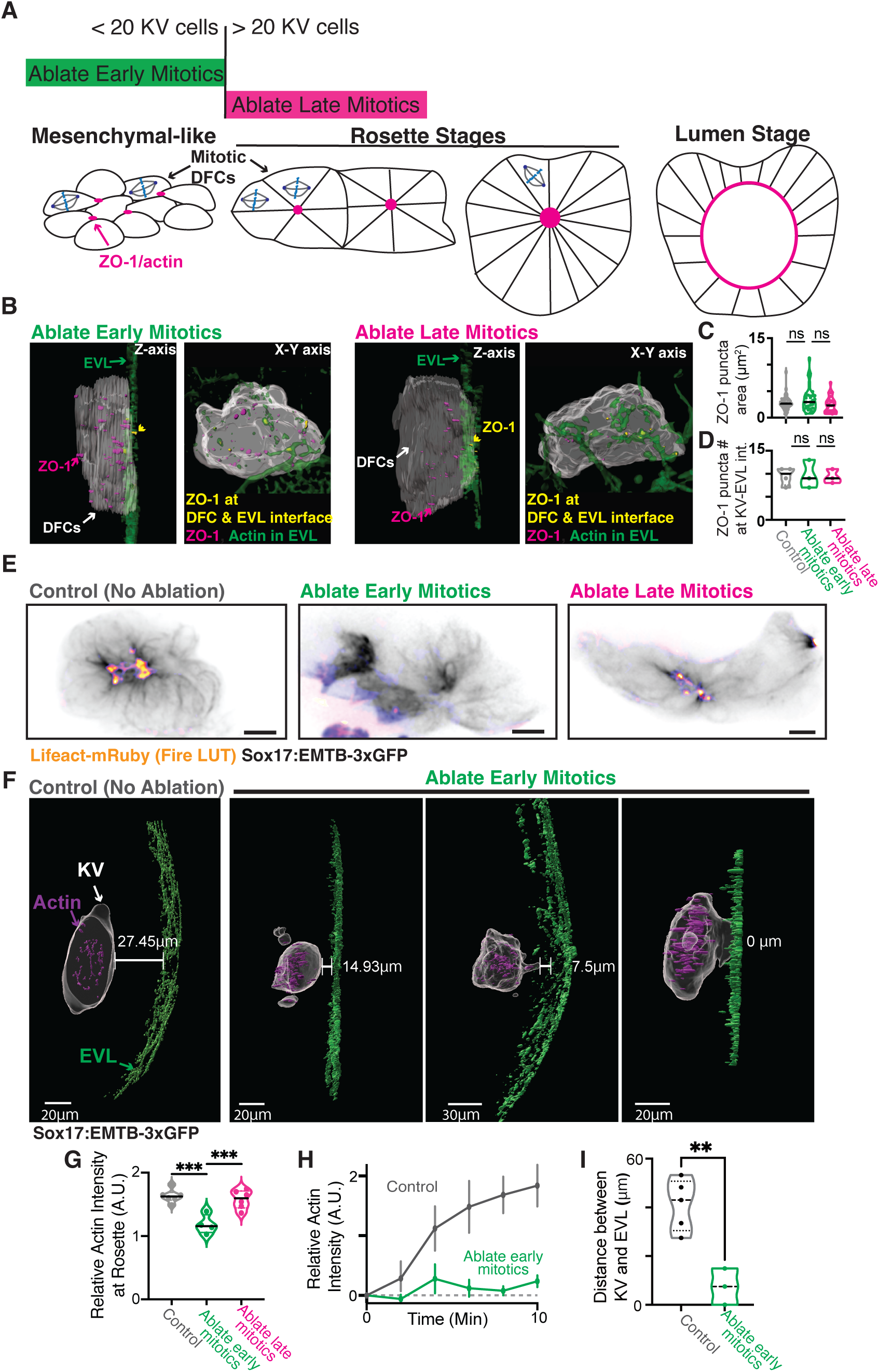
Early mitotic events drive DFC rosette transition and detachment from EVL. **(A)** Model illustrating the timing of laser ablation and ZO-1/actin assembly during KV development. **(B)** Side (Z-axis) and bottom (XY-axis) views of 3D surface renderings of representative images from embryos ablated of early mitotics compared to ablated late mitotics fixed post 20 KV cells when they are initiating small rosette formation. Dorsal forerunner cells (DFCs, gray), the enveloping layer (EVL, green), ZO-1 puncta within the KV (magenta), and ZO-1 puncta at the KV–EVL junction (yellow) are shown. **(C)** Truncated violin plot showing the projection area of ZO-1 puncta at the KV–EVL junction in embryos with ablated early mitotics vs. ablated late mitotics. Median values are indicated by black lines. ns, p > 0.05 (unpaired t-test). Control (No ablation): 2 clutches, 4 embryos, 38 puncta; ablated early mitotics: 3 clutches, 3 embryos, 29 puncta; ablated late mitotics: 3 clutches, 3 embryos, 28 puncta. **(D)** Truncated violin plot showing the number of ZO-1 puncta at the KV–EVL interface in embryos with ablated early mitotics compared to ablated late mitotics. Median values shown with black line. ns, p > 0.05 (one-way ANOVA; 3 clutches for each ablation condition and 2 clutches for control, at least 3 embryos per condition). **(E)** Representative image of actin (Lifeact-mRuby, FireLUT) in the KV rosette stage (Sox17:EMTB-3xGFP, gray) of no ablation control, ablated early mitotics, and ablated late mitotics. Scale bar 10μm. **(F)** 3D surface renderings of DFCs (gray) in no-ablation control and embryos with ablated early mitotics. The distance between DFCs and the EVL (green) is highlighted by white lines; actin puncta within the KV are shown in purple. Scale bar: 20 or 30 μm. Refer to Supplemental Video 4. **(G)** Violin plot showing relative actin intensity at rosettes for no-ablation controls, embryos ablated of early mitotics and ablated of late mitotics. Median values (black line) and quartiles (dashed lines) are shown. ***P < 0.001 (one-way ANOVA with Dunnett’s multiple comparisons; n ≥ 4 rosettes). **(H)** Relative actin intensity recruited to microtubule bundles in the KV for no-ablation control embryos (gray) and early mitotic ablated embryos (green) over 10 minutes post microtubule bundle formation. Error bars represent s.e.m.; n ≥ 5 bundles. Control (No ablation): 4 clutches, 4 embryos; ablated early mitotics: 3 clutches, 3 embryos. **(I)** Truncated violin plot showing the distance between the KV and EVL as quantified in (**F**). **p< 0.005 (unpaired t-test; n ≥ 3 embryos measured per condition). Refer to Supplemental Table S1.

We next asked whether early mitotic ablation affects rosette formation and subsequent KV detachment from the EVL. To address this, we performed live imaging of Sox17:EMTB-3xGFP embryos injected with Lifeact-mRuby and followed KV development through 12 hpf under control, early mitotic ablation, and late mitotic ablation conditions. In control embryos, multiple rosette-like structures formed by ∼9 hpf, characterized by converging microtubule bundles and robust actin accumulation at rosette centers (Figure 6E). In contrast, embryos subjected to early mitotic ablation exhibited a marked reduction in actin enrichment at prospective rosette centers, whereas ablation of later mitotic events had little effect (Figure 6E). Quantification of actin intensity at rosette centers relative to the surrounding KV tissue confirmed a significant decrease in actin recruitment following early, but not late, mitotic ablation (Figure 6G).

To assess whether early mitotic divisions are required for actin accumulation at nascent rosette sites, we quantified actin recruitment over a 10-minute time course at identified microtubule bundles. In control embryos, actin intensity increased robustly at these sites, whereas embryos lacking early mitotic events exhibited significantly reduced actin accumulation at microtubule bundles associated with later or non-mitotic events (Figure 6H). We evaluated the consequences of these defects on KV positioning relative to the EVL. Segmentation of DFCs and actin at 12 hpf revealed that KVs in embryos lacking early mitotic events remained significantly closer to the EVL compared to controls, and in some cases remained partially attached (Figure 6F, 6I, Video S4). These findings suggest that early mitotic events are dispensable for the establishment of DFC–EVL attachments but are essential for the cytoskeletal remodeling required to form actin-rich rosettes and promote KV detachment from the EVL.

## DISCUSSION

Our data support a model in which cytokinesis contributes directly to tissue morphogenesis by organizing cytoskeletal architecture. Although cytokinesis is often considered the terminal phase of cell division, a growing body of work in epithelial systems has demonstrated that cytokinetic bridges and midbodies can function as spatial landmarks for polarity establishment and junctional remodeling (Dionne et al., 2015). KV development provides an in vivo context in which these principles can be examined during a developmental transition, as DFCs progress from mesenchymal-like states into a cohesive cyst of polarized, ciliated epithelial-like cells. By showing that cytokinetic bridges recruit actin and serve as focal points for rosette assembly, our findings place cytokinesis-associated structures within the broader framework of epithelial junction formation, extending their role to settings in which polarity and epithelial organization emerge progressively during development.

Our findings distinguish between the establishment of cell–cell attachments and their subsequent reorganization. DFC–EVL junctions form independently of mitotic activity, consistent with prior work showing that a subset of DFCs inherits polarity from the surface epithelium. However, these external attachments are insufficient to drive internal epithelial organization. Instead, we propose a model where early cytokinetic events generate microtubule-based scaffolds that reorganize cytoskeletal and junctional components toward DFC–DFC interfaces, providing a mechanism for converting externally derived polarity into an internally coherent epithelial-like architecture.

Our quantitative analysis suggests that ZO-1 assembly precedes actin recruitment at DFC–EVL interfaces, with actin enrichment occurring only after junctional complexes reach a critical size or intensity. This hierarchy is consistent with models in which tight junction scaffolds provide a platform for cytoskeletal stabilization (Citi et al., 2024; Mira-Osuna and Le Borgne, 2024). Notably, once cytokinetic bridges form, actin recruitment becomes tightly coupled to microtubule organization at these sites. Laser ablation experiments establish that intact cytokinetic bridges are required for robust actin accumulation, providing an argument that microtubule structures associated with cytokinesis actively organize actin-based architectures. These observations suggest that cytokinetic bridges function as transient, polarized signaling hubs that concentrate cytoskeletal regulators and junctional components. Successive division events reinforce these sites, leading to progressive rosette maturation. This iterative mechanism provides a means by which proliferative dynamics are translated into stable tissue architecture.

Our findings provide a mechanistic explanation for the stage-specific requirement for early DFC divisions previously observed (Wu et al., 2025). Early mitotic events occur when DFCs are still engaged with the EVL and before internal epithelial architecture is established. By driving rosette formation and facilitating detachment from the EVL, these early divisions initiate the morphogenetic program that ultimately enables lumen formation. Later divisions, by contrast, occur after rosettes and internal junctional networks are established and therefore do not play a comparable organizational role and have been shown to be dispensable (Wu et al., 2025). This temporal distinction highlights how the developmental context of cell division, not simply division number, determines its morphogenetic impact.

In summary, this study positions cytokinesis as an instructive morphogenetic event that drives cytoskeletal reorganization, rosette formation, and epithelialization during left–right organizer development. These findings expand the functional repertoire of cytokinesis beyond cell division and provide new insight into how complex tissue architectures can arise de novo during embryonic development.

## ACKNOWLEDGEMENTS

This work was supported by National Institute of General Medical Sciences (R01GM-127621, R01GM-130874 and R35GM158119 to H.H.). We thank NSF MicroFFABS REU program at Syracuse University for their support of M.A.A..

## AUTHOR CONTRIBUTIONS

Conceptualization: Y.W., Y.L., H.H.; Data Curation: Y.W., Y.L., M.A.A., C.T.; Formal analysis: Y.W., Y.L., M.A.A., C.T.; Funding Acquisition: H.H.; Investigation: Y.W., Y.L., M.A.A., C.T.; Methodology: Y.W., Y.L., M.A.A., C.T.; Project administration: Y.W., Y.L., H.H.; Resources: Y.W., Y.L., M.A.A., C.T.; Supervision: H.H.; Validation: Y.W., Y.L., M.A.A., C.T., H.H.; Visualizaiton: Y.W., Y.L., M.A.A., C.T., H.H.; Writing-original draft: H.H.; Writing-review & editing: Y.W., Y.L., M.A.A., C.T., H.H.

---

**Video S1**: *DFC first attach to the EVL through ZO-1, then detach during rosette formation, KV rounding, and lumenogenesis.* 3D surface renderings of representative embryos showing dorsal forerunner cells (DFCs, gray), enveloping layer (EVL, green), ZO-1 visualized in magenta. Related to Figure 2B.

**Video S2**: *ZO-1 site assembly precedes actin; ZO-1 and actin then accumulate together at DFC-EVL interfaces.* Live confocal video showing KV cell nucleus (Sox17:H2B-mCherry, mid gray scale, magenta in merged), ZO-1 (Tgki(Tjp1atdTomato), mid gray scale, magenta in merged), and actin (Lifeact-mEmerald, left gray scale, cyan in merged). Scale bar: 10 µm. Related to Figure 3E.

**Video S3**: *Once ZO-1-EVL attachments form, cytokinetic events converge and precede rosette formation*. Time lapse of actin (Lifeact-mRuby, fire-LUT) and KV MTs (Sox17:EMTB-3xGFP, shown in inverted gray) during KV rosette formation. Scale bar, 10 μm. Related to Figure 4B-C.

**Video S4**: *Early mitotic events drive DFC rosette transition and detachment from EVL.* 3D surface renderings of DFCs (gray) in no-ablation control and when early mitotics were ablated in embryos, KV Boundary (Sox17:EMTB-3xGFP, inverted gray), Actin shown in purple, EVL shown in green. Scale bar, 20μm. Related to Figure 6F.

## MATERIALS AND METHODS

### Fish lines

Zebrafish lines were maintained in accordance with protocols approved by the Institutional Animal Care Committee of Syracuse University (IACUC Protocol #18-006). Embryos were raised at 28.5°C and staged (as described by (Kimmel et al., 1995)). Wild-type and/or transgenic zebrafish lines used for live imaging and immunohistochemistry are listed in Resource Table (Supplemental Table S2).

### Plasmids and mRNA for injection experiments

Plasmids were generated using Gibson cloning methods (NEBuilder HiFi DNA assembly Cloning Kit) and maxi-prepped. mRNA was made using the mMESSAGE mMACHINE™ SP6 transcription kit. See Resource Table (Supplemental Table S2) for a list of plasmid constructs and mRNAs used. Injections of one-cell-staged embryos were performed as described in (Aljiboury et al., 2021).

### Immunofluorescence

Immunostaining for ZO-1, Actin and GFP, Tg(Sox17:EMTB-3xGFP) transgenic zebrafish embryos were used and were fixed at 6, 9, 10, and 12 hpf using 4% PFA with 0.5% Triton X-100 (dissolved in PBS) overnight at 4°C. Embryos were dechorionated after washing with PBST (PBS with 0.1% Triton X-100) three times. Embryos were blocked in wash solution (1% DMSO, 1% bovine serum albumin, 0.1% Triton X-100) for 1 h at room temperature with gentle agitation. Primary antibody incubation (diluted in wash solution) was carried out overnight at 4°C. Primary antibodies used were: anti-GFP (chicken) (1:300; GeneTex, GTX13970; AB_371416), and anti-ZO-1 (mouse) (1:400; Invitrogen, 339194); see Resource Table (Supplemental Table S2). Embryos were then washed and incubated with secondary antibodies for 2-4 h at room temperature or overnight at 4°C. Secondary antibodies used were: Alexa Fluor anti-chicken 488 (1:300; Fisher Scientific, A11039), and Phalloidin (1:1000; Cell Signaling Technology, 8940S). Embryos were stained with DAPI (1 μg/ml) to label nuclei after three washes with wash solution. Embryos were mounted in 1.2% agarose after washing with PBS. See Resource Table (Supplemental Table S2).

### Imaging

Fixed or live dechorionated Tg(Sox17:EMTB-3xGFP) or Tg(Sox17:EMTB-3xGFP; ZO-1-tdTomato) embryos are embedded in low-melting 1.2% agarose (Supplemental Table S2) with the KV positioned at the bottom of a #1.5 glass-bottom MatTek plate (Supplemental Table S2) and imaged. Zebrafish embryos were imaged using either a Leica DMi8 inverted microscope equipped with a X-light V2 confocal unit spinning disk equipped with a Visitron VisiFRAP-DC photokinetics system attached to 405 and 355 nm lasers, a Dragonfly-620 SR spinning disk confocal, or a Leica SP8 laser scanner confocal microscope (LSCM). The Leica DMi8 is equipped with a Lumencor SPECTRA X, Photometrics Prime-95B sCMOS camera and 89 North-LDi laser launch. VisiView software was used to acquire images. Optics used with this unit were: HC PL APO ×40/1.10W CORR CS2 0.65 water immersion objective, HC PL APO×40/0.95 NA CORR dry objective and HCX PL APO ×63/1.40-0.06 NA oil objective. The SP8 laser scanning confocal microscope is equipped with an HC PL APO ×20/0.75 IMM CORR CS2 objective, HC PL APO ×40/1.10 W CORR CS2 0.65 water objective and HC PL APO ×63/1.3 Glyc CORR CS2 glycerol objective. LAS-X software was used to acquire images. The Dragonfly-520 SR spinning disk confocal is equipped with MicroPoint Gen 4 photostimulation tool, ZL41 sCMOS camera and iXON EMCCD camera for acquisition. Optics used with units is LD LCI Plan-Apochromat 40x/1.2 Imm Corr DIC M27 water immersion objective. Fusion 2.3 was used to acquire images. A Leica M165 FC stereomicroscope equipped with DFC 9000 GT sCMOS camera was used for staging and phenotypic analysis of zebrafish embryos.

### Laser ablation

For ablation of mitotic events, dechorionated live Tg(Sox17:EMTB-3xGFP) embryos were mounted and imaged on a spinning-disk confocal microscope equipped with a VisiView kinetics unit. Imaging was initiated at either 6 hpf (to ablate early mitotic events, when KV cell number was <20) or 7 hpf (to ablate later mitotic events). Time-lapse image stacks were acquired using 470 nm and 555 nm lasers to visualize EMTB-3xGFP and mRuby-Lifeact, respectively, with z-stacks collected at 2 μm intervals every 2 min and continued until 12 hpf. Mitotic events within the KV were identified by spindle formation and chromosome alignment and were ablated using a 355 nm laser at 65% power. Ablation was performed by applying the laser to a defined region of interest (ROI) encompassing the mitotic cell for two cycles of 50 ms per pixel.

For ablation of cytokinetic microtubule bundles, embryos were imaged under the same conditions until the formation of cytokinetic bridge–derived microtubule bundles were detected. Targeted line ablations (∼2 μm in length) were applied to microtubule bundles in one daughter cell using a 355 nm laser at 60% power, while the opposing daughter cell was left intact as an internal control. The laser was applied within a defined ROI centered on the microtubule bundle for two cycles of 50 ms per pixel. All ablations were performed prior to robust actin accumulation at the targeted site.

### Image and data analysis

Images were processed using Fiji/ImageJ and Imaris (Bitplane). Graphs and statistical analysis were produced using Prism 9 software. Movies were created using Fiji/ImageJ or IMARIS. All time-lapse movie projections are registered using Fiji.

### Relative actin intensity

Relative actin intensity at rosette centers was quantified using Fiji/ImageJ. A fixed 6 μm × 6 μm rectangular ROI was placed over the center of each rosette, as defined by converging microtubule bundles and enriched Lifeact signal. Local background intensity was measured using an identically sized ROI positioned in an adjacent non-rosette region within the same KV. Rosette actin intensity was calculated by dividing the mean fluorescence intensity within the rosette ROI by the mean fluorescence intensity of the local background ROI.

For analysis of actin recruitment at cytokinetic microtubule bundles, a 3 μm × 3 μm ROI was placed at the tip of each microtubule bundle, and Lifeact-mRuby fluorescence was measured over a 10-minute period following laser ablation. At each time point, relative actin intensity was calculated by subtracting the minimum fluorescence intensity measured across the time series from the mean ROI intensity and normalizing this value to the intensity at 0 minutes post-ablation.

### Quantification of actin intensity relative to ZO-1 puncta

Actin and ZO-1 fluorescence intensities at individual ZO-1 puncta were quantified using Fiji/ImageJ. Each punctum was manually outlined using a freehand ROI. Local background fluorescence was measured by placing an identically shaped ROI in an adjacent region lacking detectable puncta. For each punctum, background-subtracted mean fluorescence intensities for ZO-1 and actin were calculated. These values were then normalized to the average fluorescence intensity measured across all puncta within the same embryo for ZO-1 and actin, respectively.

### Quantification of ZO-1 puncta area

Three-dimensional (3D) surface rendering and quantitative analysis of ZO-1 puncta were performed using Imaris (Oxford Instruments) to assess puncta located within the KV or at the KV–EVL interface. KV and EVL surfaces were generated independently using the Surfaces module. The EVL was first segmented using either the actin or ZO-1 fluorescence channel to generate an accurate 3D surface representation. The KV was then isolated and rendered as a separate surface. KV and EVL surfaces were pseudo colored to facilitate visual distinction during analysis. Individual ZO-1 puncta were manually identified, isolated, and masked. Puncta were classified as KV–EVL junctional if they were in direct contact with both the KV and EVL surfaces. Following masking, the volume of each ZO-1 punctum was quantified using the Imaris volume measurement function.

### Surface rendering and measurement of KV–EVL distance

3D surface rendering and quantitative distance measurements were performed using Imaris (Oxford Instruments) to assess the spatial relationship between the KV and the EVL in zebrafish embryos. KV and EVL surfaces were generated independently using the Surfaces module. The EVL was first segmented using either the actin or ZO-1 fluorescence channel to generate an accurate 3D surface representation of the epithelial layer. The KV was then isolated and rendered as a distinct surface based on EMTB-3xGFP labeling. Following segmentation, KV and EVL surfaces were pseudocolored to facilitate visual distinction during analysis. The shortest distance between the KV boundary and the EVL surface was quantified using the “Shortest Distance to Surface” function within the Imaris Statistics module. Distance measurements were obtained for each embryo and compiled for downstream statistical analysis.

### Statistical analysis

Prism 9 software was used for all graph preparations, which include all individual data points across embryos and as noted in legends. Unpaired, two-tailed t-tests for pairwise numerical differences in mean and one-way ANOVA were performed when comparing greater than two groups using Prism 9 software. Pearson’s correlation was performed on investigating correlation between two sets of data using Prism 9 software. P-values are represented as: ****P<0.0001, ***P<0.001, **P<0.01, *P<0.05; ns, not significant. All n values and statistical analysis information for each figure are included in Statistics Table (Supplemental Table S1).

---

**Table S1.**
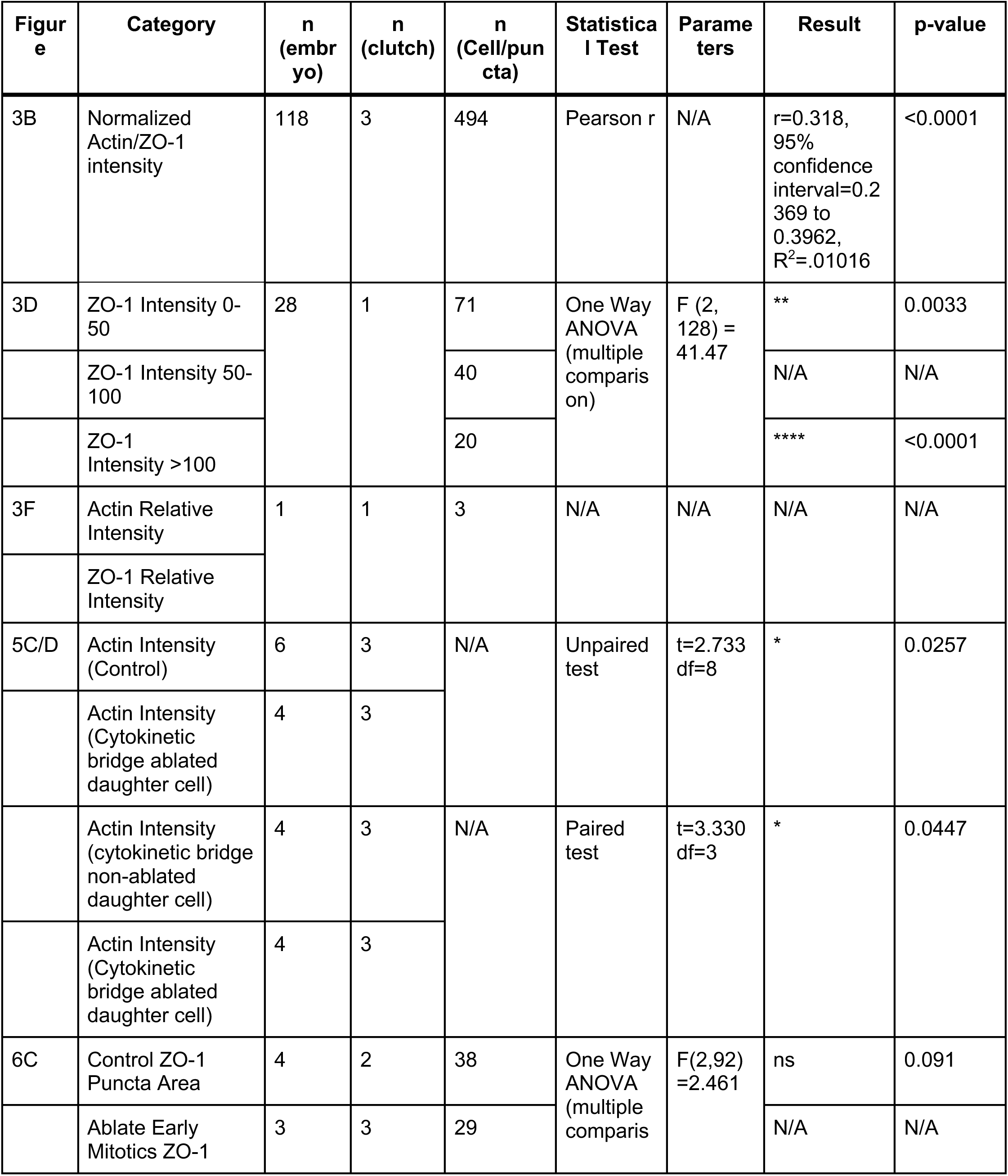
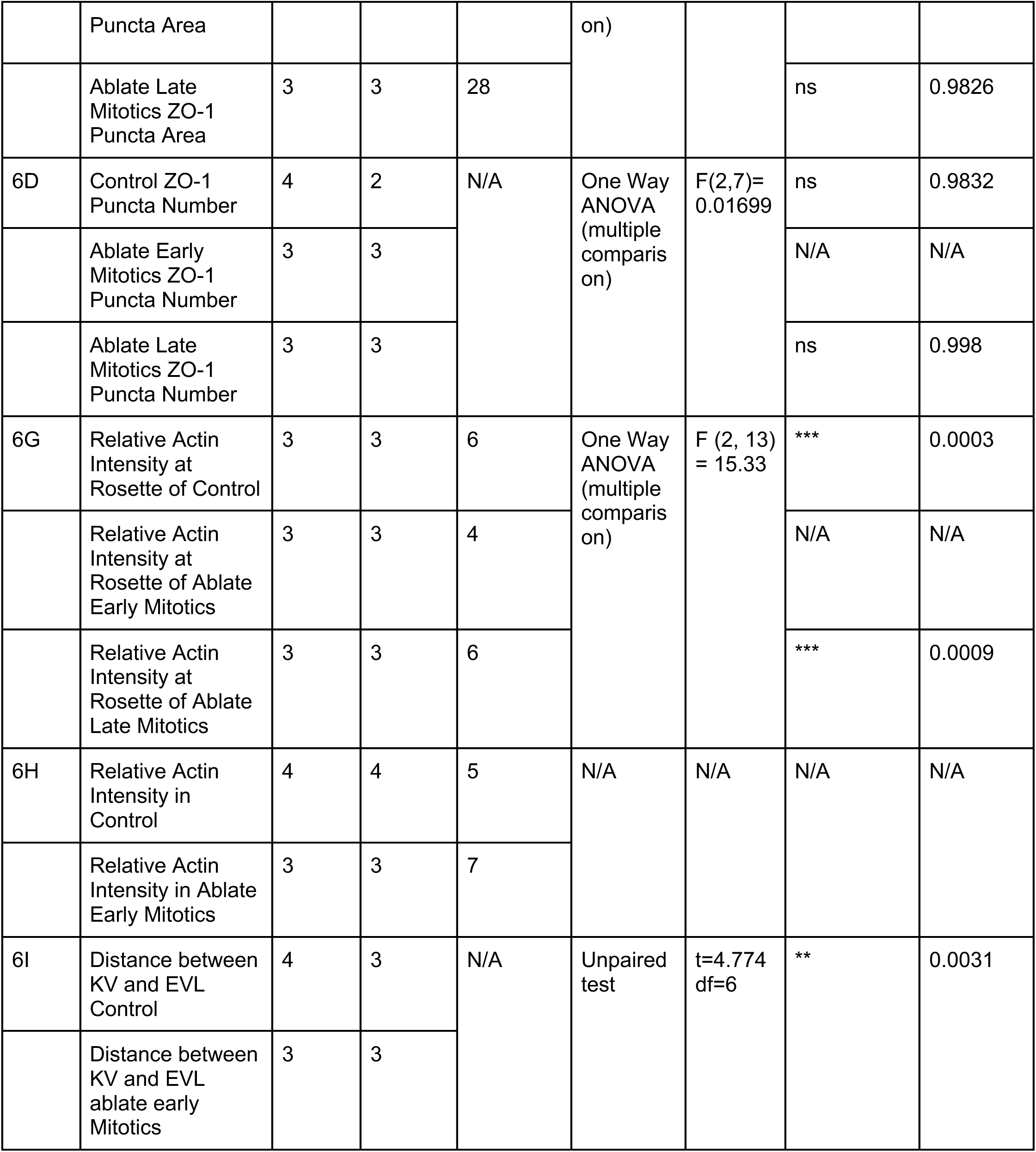
Detailed statistical analysis of results reported in this study.

**Table S2.** Reagents and Tools Table.

| Reagent/Resource | Reference or Source | Identifier or Catalog Number |
| --- | --- | --- |
| **Experimental Models** |  |  |
| Zebrafish | Zebrafish International Resource Center | AB-Wildtype |
| Zebrafish | Michel Bagnat lab | Tgki(Tjp1atdTomato) |
| Zebrafish | Heidi Hehnly lab | Tg(Sox17:EMTB-3xGFP) |
| Zebrafish | Heidi Hehnly lab | Tg(Sox17:H2B-mCherry) |
| Zebrafish | David M Tobin Lab | cdh1-mlanYFP |
| **DNA** |  |  |
| Plasmid:pCS2-Lifeact-mEmerald | Addgene | Plasmid #252242 |
| Plasmid: pCS2-Lifeact-mRuby | Addgene | Plasmid #194339 |
| **Antibodies** |  |  |
| Anti-GFP (Chicken) | GeneTex | GTX13970: AB_371416 |
| ZO-1 Monoclonal Antibody (ZO1-1A12), Alexa Fluor 594, Alexa Fluor® 594, Mouse | Invitrogen | 339194 |
| Alexa Fluor Anti-Chicken 488 | Fisher scientific | A11039 |
| Alexa Fluor® 647 Phalloidin | Cell Signaling Technology | 8940S |
| **Oligonucleotides and other sequence-based reagents** |  |  |
| Lifeact-mRuby | Heidi Hehnly lab | Plasmid: pCS2-Lifeact-mRuby |
| Lifeact-mEmerald | Heidi Hehnly lab | Plasmid: pCS2-Lifeact-mEmerald |
| **Chemicals, Enzymes and other reagents** |  |  |
| DAPI | Sigma Aldrich | D9542-10mg |
| Agarose | Thermo Fisher | 16520100 |
| BSA | Fisher Scientific | BP1600-100 |
| QIAGEN Plasmid Midi Kit | Qiagen | 12143 |
| BIO BASIC Maxi Prep Kit | BIO BASIC | 9K-0060023 |
| Dimethylsulphoxide | Fisher Scientific | BP231-100 |
| Paraformaldehyde | Fisher Scientific | O4042-500 |
| Phosphate Buffered Saline | Fisher Scientific | 10010023 |
| Triton X-100 | Fisher Scientific | BP151500 |
| Tween 20 | Thermo Fisher | BP337500 |
| Sodium Chloride | Fisher Scientific | BP358 |
| NEBuilder HiFi DNA assembly Cloning Kit | New England BioLabs | E5520S |
| mMESSAGE mMACHINETMSP6 | Invitrogen | AM1340 |
| **Software** |  |  |
| ImageJ/FIJI | NIH and Laboratory for<br>Optical and Computational<br>Instrumentation | https://imagej.net/Fiji |
| IMARIS, Bitplane | Oxford Instruments | https://imaris.oxinst.com/ |
| PRISM9 | GraphPad | https://www.graphpad.com/scientific-software/prism/ |
| LAS-X Software | Leica Microsystems | https://www.leica-microsystems.com/products/microscope-software/p/leica-las-x-ls/ |
| VisiView | Visitron | https://www.visitron.de/products/visiviewr-software.html |
| Fusion | Oxford Instruments | https://fusion-benchtop-software-guide.scrollhelp.site/fusionum/ |
| **Other** | N/A | N/A |

## References

Addi, C., Bai, J. and Echard, A. (2018). Actin, microtubule, septin and ESCRT filament remodeling during late steps of cytokinesis. Curr. Opin. Cell Biol. 50, 27–34.

Aljiboury, A. A., Mujcic, A., Cammerino, T., Rathbun, L. I. and Hehnly, H. (2021). Imaging the early zebrafish embryo centrosomes following injection of small-molecule inhibitors to understand spindle formation. STAR Protoc. 2, 100293.

Amack, J. D. (2021). Cellular dynamics of EMT: lessons from live in vivo imaging of embryonic development. Cell Commun. Signal. 19, 79.

Amack, J. D., Wang, X. and Yost, H. J. (2007). Two T-box genes play independent and cooperative roles to regulate morphogenesis of ciliated Kupffer’s vesicle in zebrafish. Dev. Biol. 310, 196–210.

Babb, S. G. and Marrs, J. A. (2004). E-cadherin regulates cell movements and tissue formation in early zebrafish embryos. Dev. Dyn. 230, 263–277.

Chartier, N. T., Salazar Ospina, D. P., Benkemoun, L., Mayer, M., Grill, S. W., Maddox, A. S. and Labbé, J. C. (2011). PAR-4/LKB1 mobilizes nonmuscle myosin through anillin to regulate C. elegans embryonic polarization and cytokinesis. Current Biology 21, 259–269.

Citi, S., Fromm, M., Furuse, M., González-Mariscal, L., Nusrat, A., Tsukita, S. and Turner, J. R. (2024). A short guide to the tight junction. J. Cell Sci. 137,.

Cronan, M. R. and Tobin, D. M. (2019). Endogenous Tagging at the cdh1 Locus for Live Visualization of E-Cadherin Dynamics. Zebrafish 16, 324–325.

Dambournet, D., MacHicoane, M., Chesneau, L., Sachse, M., Rocancourt, M., El Marjou, A., Formstecher, E., Salomon, R., Goud, B. and Echard, A. (2011). Rab35 GTPase and OCRL phosphatase remodel lipids and F-actin for successful cytokinesis. Nat. Cell Biol. 13, 981–988.

Daniel, E., Daudé, M., Kolotuev, I., Charish, K., Auld, V. and Le Borgne, R. (2018). Coordination of Septate Junctions Assembly and Completion of Cytokinesis in Proliferative Epithelial Tissues. Current Biology 28, 1380–1391.e4.

Dasgupta, A. and Amack, J. D. (2016). Cilia in vertebrate left-right patterning. Philos. Trans. R. Soc. Lond. B Biol. Sci. 371,.

Dasgupta, A., Merkel, M., Clark, M. J., Jacob, A. E., Dawson, J. E., Manning, M. L. and Amack, J. D. (2018). Cell volume changes contribute to epithelial morphogenesis in zebrafish Kupffer’s vesicle. Elife 7,.

di Pietro, F., Osswald, M., De las Heras, J. M., Cristo, I., López-Gay, J., Wang, Z., Pelletier, S., Gaugué, I., Leroy, A., Martin, C., et al. (2023). Systematic analysis of RhoGEF/GAP localizations uncovers regulators of mechanosensing and junction formation during epithelial cell division. Current Biology 33, 858–874.e7.

Dionne, L. K., Wang, X. J. and Prekeris, R. (2015). Midbody: From cellular junk to regulator of cell polarity and cell fate. Curr. Opin. Cell Biol. 35, 51–58.

Erdemci-Tandogan, G., Clark, M. J., Amack, J. D. and Manning, M. L. (2018). Tissue Flow Induces Cell Shape Changes During Organogenesis. Biophys. J. 115, 2259–2270.

Essner, J. J., Amack, J. D., Nyholm, M. K., Harris, E. B. and Yost, H. J. (2005). Kupffer’s vesicle is a ciliated organ of asymmetry in the zebrafish embryo that initiates left-right development of the brain, heart and gut. Development 132, 1247–1260.

Frisca, F., Colquhoun, D., Goldshmit, Y., Änkö, M. L., Pébay, A. and Kaslin, J. (2016). Role of ectonucleotide pyrophosphatase/phosphodiesterase 2 in the midline axis formation of zebrafish. Sci. Rep. 6,.

Gredler, M. L. and Zallen, J. A. (2023). Multicellular rosettes link mesenchymal-epithelial transition to radial intercalation in the mouse axial mesoderm. Dev. Cell 58, 933–950.e5.

Grimes, D. T. and Burdine, R. D. (2017). Left–Right Patterning: Breaking Symmetry to Asymmetric Morphogenesis. Trends in Genetics 33, 616–628.

Guillot, C. and Lecuit, T. (2013). Mechanics of epithelial tissue homeostasis and morphogenesis. Science 340, 1185–1189.

Harding, M. J., McGraw, H. F. and Nechiporuk, A. (2014). The roles and regulation of multicellular rosette structures during morphogenesis. Development 141, 2549–2558.

Herszterg, S., Leibfried, A., Bosveld, F., Martin, C. and Bellaiche, Y. (2013). Interplay between the Dividing Cell and Its Neighbors Regulates Adherens Junction Formation during Cytokinesis in Epithelial Tissue. Dev. Cell 24, 256–270.

Hirokawa, N., Tanaka, Y., Okada, Y. and Takeda, S. (2006). Nodal Flow and the Generation of Left-Right Asymmetry. Cell 125, 33–45.

Hirokawa, N., Tanaka, Y. and Okada, Y. (2009). Left–Right Determination: Involvement of Molecular Motor KIF3, Cilia, and Nodal Flow. Cold Spring Harb. Perspect. Biol. 1,.

Kane, D. A., Hammerschmidt, M., Mullins, M. C., Maischein, H. M., Brand, M., Van Eeden, F. J. M., Furutani-Seiki, M., Granato, M., Haffter, P., Heisenberg, C. P., et al. (1996). The zebrafish epiboly mutants. Development 123, 47–55.

Kimmel, C. B., Ballard, W. W., Kimmel, S. R., Ullmann, B. and Schilling, T. F. (1995). Stages of embryonic development of the zebrafish. Developmental dynamics : an official public 203, 253–310.

Kramer-Zucker, A. G., Olale, F., Haycraft, C. J., Yoder, B. K., Schier, A. F. and Drummond, I. A. (2005). Cilia-driven fluid flow in the zebrafish pronephros, brain and Kupffer’s vesicle is required for normal organogenesis. Development 132, 1907–21.

Krishnan, N., Swoger, M., Rathbun, L. I., Fioramonti, P. J., Freshour, J., Bates, M., Patteson, A. E. and Hehnly, H. (2022). Rab11 endosomes and Pericentrin coordinate centrosome movement during pre-abscission in vivo. Life Sci. Alliance 5, 1–15.

Lee, J. D. and Anderson, K. V. (2008). Morphogenesis of the node and notochord: the cellular basis for the establishment and maintenance of left-right asymmetry in the mouse. Dev. Dyn. 237, 3464–3476.

Levic, D. S., Yamaguchi, N., Wang, S., Knaut, H. and Bagnat, M. (2021). Knock-in tagging in zebrafish facilitated by insertion into non-coding regions. Development 148,.

Lopes, S. S., Lourenço, R., Pacheco, L., Moreno, N., Kreiling, J. and Saúde, L. (2010). Notch signalling regulates left-right asymmetry through ciliary length control. Development 137, 3625–3632.

Mangan, A. J., Sietsema, D. V, Li, D., Moore, J. K., Citi, S. and Prekeris, R. (2016). Cingulin and actin mediate midbody-dependent apical lumen formation during polarization of epithelial cells. Nat. Commun. 7, 12426.

Manna, R. K., Retzlaff, E. M., Hinman, A. M., Lan, Y., Abdel-Razek, O., Bates, M., Hehnly, H., Amack, J. D. and Lisa Manning, M. (2025). Dynamic forces drive cell and organ morphology changes during embryonic development. Proc. Natl. Acad. Sci. U. S. A. 122,.

Mira-Osuna, M. and Le Borgne, R. (2024). Assembly, dynamics and remodeling of epithelial cell junctions throughout development. Development (Cambridge) 151,.

Navis, A., Marjoram, L. and Bagnat, M. (2013). Cftr controls lumen expansion and function of Kupffer’s vesicle in zebrafish. Development 140, 1703–12.

Okabe, N., Xu, B. and Burdine, R. D. (2008). Fluid dynamics in zebrafish Kupffer’s vesicle. Dev. Dyn. 237, 3602–3612.

Oteíza, P., Köppen, M., Concha, M. L. and Heisenberg, C. P. (2008). Origin and shaping of the laterality organ in zebrafish. Development 135, 2807–2813.

Pinheiro, D. and Bellaïche, Y. (2018). Mechanical Force-Driven Adherens Junction Remodeling and Epithelial Dynamics. Dev. Cell 47, 3–19.

Pinheiro, D., Hannezo, E., Herszterg, S., Bosveld, F., Gaugue, I., Balakireva, M., Wang, Z., Cristo, I., Rigaud, S. U., Markova, O., et al. (2017). Transmission of cytokinesis forces via E-cadherin dilution and actomyosin flows. Nature 2017 545:7652 545, 103–107.

Rathbun, L. I., Colicino, E. G., Manikas, J., O’Connell, J., Krishnan, N., Reilly, N. S., Coyne, S., Erdemci-Tandogan, G., Garrastegui, A., Freshour, J., et al. (2020). Cytokinetic bridge triggers de novo lumen formation in vivo. Nat. Commun. 11, 1–12.

Sampaio, P., Ferreira, R. R., Guerrero, A., Pintado, P., Tavares, B., Amaro, J., Smith, A. A., Montenegro-Johnson, T., Smith, D. J. and Lopes, S. S. (2014). Left-right organizer flow dynamics: How much cilia activity reliably yields laterality? Dev. Cell 29, 716–728.

Sanematsu, P. C., Erdemci-Tandogan, G., Patel, H., Retzlaff, E. M., Amack, J. D. and Manning, M. L. (2021). 3D viscoelastic drag forces contribute to cell shape changes during organogenesis in the zebrafish embryo. Cells and Development.

Schlüter, M. A., Pfarr, C. S., Pieczynski, J., Whiteman, E. L., Hurd, T. W., Fan, S., Liu, C.-J. and Margolis, B. (2009). Trafficking of Crumbs3 during cytokinesis is crucial for lumen formation. Mol. Biol. Cell 20, 4652–63.

Shimizu, T., Yabe, T., Muraoka, O., Yonemura, S., Aramaki, S., Hatta, K., Bae, Y. K., Nojima, H. and Hibi, M. (2005). E-cadherin is required for gastrulation cell movements in zebrafish. Mech. Dev. 122, 747–763.

Sugioka, K. (2022). Symmetry-breaking of animal cytokinesis. Semin. Cell Dev. Biol. 127, 100–109.

Tay, H. G., Schulze, S. K., Compagnon, J., Foley, F. C., Heisenberg, C. P., Yost, H. J., Abdelilah-Seyfried, S. and Amack, J. D. (2013). Lethal giant larvae 2 regulates development of the ciliated organ Kupffer’s vesicle. Development (Cambridge) 140, 1550–1559.

Wang, G., Cadwallader, A. B., Jang, D. S., Tsang, M., Yost, H. J. and Amack, J. D. (2011). The Rho kinase rock2b establishes anteroposterior asymmetry of the ciliated Kupffer’s vesicle in zebrafish. Development 138, 45–54.

Wang, G., Manning, M. L. and Amack, J. D. (2012). Regional cell shape changes control form and function of Kupffer’s vesicle in the zebrafish embryo. Dev. Biol. 370, 52–62.

Wang, T., Yanger, K., Stanger, B. Z., Cassio, D. and Bi, E. (2014). Cytokinesis defines a spatial landmark for hepatocyte polarization and apical lumen formation. J. Cell Sci. 127, 2483–92.

Wells, J. R., Padua, M. B. and Ware, S. M. (2022). The genetic landscape of cardiovascular left-right patterning defects. Curr. Opin. Genet. Dev. 75,.

Wu, Y., Lan, Y., Ononiwu, F., Poole, A., Rasmussen, K., Silva, J. Da, Shamil, A. W., Jao, L.-E. and Hehnly, H. (2025). Specific mitotic events drive left-right organizer development. Development.

Zhang, J., Jiang, Z., Liu, X. and Meng, A. (2016). Eph/ephrin signaling maintains the boundary of dorsal forerunner cell cluster during morphogenesis of the zebrafish embryonic left-right organizer. Development 143, 2603–2615.

